# CHK2 inhibition provides a strategy to suppress hematological toxicity from PARP inhibitors

**DOI:** 10.1101/2020.07.28.222927

**Authors:** Zhen Xu, Cassandra J Vandenberg, Elizabeth Lieschke, Clare L Scott, Ian J Majewski

## Abstract

Cancer patients treated with poly (ADP-ribose) polymerase inhibitors (PARPi) experience various side effects, with hematological toxicity being most common. Short term treatment of mice with olaparib resulted in depletion of reticulocytes, B cell progenitors and immature thymocytes, whereas longer treatment induced broader myelosuppression. We performed a CRISPR/Cas9 screen targeting DNA repair genes to identify strategies to suppress hematological toxicity. The screen revealed that sgRNAs targeting the serine/threonine kinase CHK2 were enriched following olaparib treatment. Genetic or pharmacological inhibition of CHK2 blunted PARPi response in lymphoid and myeloid cell lines, and in primary pre-B/pro-B cells. Using a Cas9 base editor, we found that blocking CHK2-mediated phosphorylation of p53 also impaired olaparib response. Our results identify the p53 pathway as a major determinant of the acute response to PARPi in normal blood cells and demonstrate that targeting CHK2 can short-circuit this response. Cotreatment with a CHK2 inhibitor did not antagonise olaparib response in ovarian cancer cells. Selective inhibition of CHK2 may spare blood cells from the toxic influence of PARPi and broaden the utility of these drugs.

## Introduction

Genome instability is a hallmark of cancer (1), and understanding precisely how DNA repair pathways are compromised can help to guide treatment. Employing a therapy to target cells with a specific DNA repair defect is attractive, because of the potential to spare normal cells. To date this concept is most advanced in cancers with defective homologous recombination (HR), where the ability to accurately repair double-strand DNA breaks (DSBs) is compromised. BRCA1 and BRCA2 are key players in HR (2) and germline mutations in these genes confer a considerable lifetime risk of cancer, particularly breast, ovarian, prostate and pancreatic cancer (2, 3). BRCA-mutant cancers show classical signs of defective DNA repair - they accumulate high mutation burdens, including a specific pattern of substitutions and rearrangements (4-6). As expected, BRCA-deficient cancers are more sensitive to agents that induce DSBs, including platinum-based chemotherapies that induce inter-strand cross-links (7). However, these agents are broadly toxic, leading to the search for ways to target cancers with HR defects more selectively.

Another way to approach treating HR deficient cancers is to attack the remaining DNA repair pathways. Pursuit of this strategy resulted in the discovery that cells with HR defects are remarkably sensitive to drugs that target poly(ADP-ribose) polymerase (PARP), a key DNA damage sensor that is essential for the repair of single strand DNA breaks (8, 9). Targeting PARP creates multiple stresses for the cell, including a near complete breakdown in base excision repair and trapped PARP complexes that trigger DSBs. The connection between BRCA and PARP is held up as the first successful example of a therapeutic strategy based on the concept of synthetic lethality (10). Multiple PARP inhibitors (PARPi), including olaparib, rucaparib, niraparib and talazoparib, are now approved to treat patients with germline BRCA mutant breast, ovarian and pancreatic cancers (2, 11, 12). Their role in BRCA-mutant ovarian cancer, for example, has been enshrined with multiple reimbursement approvals in the maintenance therapy setting, and most recently, in first-line disease, regardless of BRCA status (12-15)

It is important to remember that PARP1 and PARP2 sit at the front line of the DNA damage response and that PARPi cause major disruptions to normal DNA repair processes. PARPi are associated with significant on-target side effects, which can be difficult to manage for some patients. The most frequent adverse events include nausea, fatigue and anemia (grade ≥3 in 17% of patients (National Cancer Institute (NCI) Common Terminology Criteria for Adverse Events (CTCAE; version 4.0))) (16). Indeed, anemia (18% grade 3) was the most common grade 3 adverse event in SOLO2, which assessed olaparib in BRCA mutated ovarian cancer, followed by neutropenia (4% grade 3), and these were the most common side effects that resulted in discontinuation of treatment (17). These toxic impacts are not surprising given the high turnover in the blood cell compartment, and the vital roles of PARP1 and PARP2 across diverse biological processes (18). Genetic studies have implicated PARP2 as being particularly important in the hematopoietic system, and extravascular hemolytic anemia and T cell defects have been reported in PARP-2 knockout mice (19, 20). As hematopoietic toxicity was commonly seen with PARPi treatment, we decided to assess the acute toxicity of olaparib in the murine hematopoietic system and use this system to identify synthetic viability interactions to negate these effects.

Based on a CRISPR/Cas9 screen performed in a pro-B/pre-B cell line, we found that inhibition of checkpoint kinase 2 (CHK2, or CHEK2) is a promising approach to achieve synthetic viability for blood cells treated with PARPi. CHK2 is a serine/threonine kinase that has a critical role in signalling in response to DNA damage. Following activation by ATM, CHK2 phosphorylates and stabilises the tumor suppressor p53, which then coordinates the cellular response to DNA damage (21-23). PARPi treatment rapidly induces p53 protein in B cell lines and primary pro-B/pre-B cells. We used CRISPR base editing to assess how phosphorylation on key serine and threonine residues contributes to p53 activation. The idea that p53 is central to the toxic impact of PARPi therapy on the hematopoietic compartment is notable, because p53 is disabled nearly universally in the cancers where PARPi are applied, particularly in ovarian and breast cancers (24, 25). This suggests co-targeting CHK2 could alleviate some side effects of PARPi in the blood, without a detrimental impact on the efficacy of these agents against the cancer. Others have suggested co-targeting CHK2 to boost the activity of PARPi (26, 27), but our findings point to a distinct rationale, more in keeping with recent efforts to harness CHK2 inhibitors to limit the toxic influence of radiation (28, 29) or cytotoxic chemotherapy (30).

## Results

### PARPi treatment reduced reticulocyte and BM immature B cell counts in mice

To assess the impact of PARPi on a normal hematopoietic system, we treated C57BL/6 mice with olaparib for 4 days and monitored mature blood cells and progenitors. After 4 days, there was no significant drop in red blood count (RBC) (Figure 1A), white blood count (WBC), hemoglobin (HGB) or platelets (PLT) (Supplementary Figure 1A-C). However, the number of reticulocytes dropped significantly (Figure 1B), indicating greater sensitivity in immature red blood cells. There was no decrease in the total cellularity of the bone marrow, lymph nodes or the spleen (Supplementary Figure 1D-F), but the number of thymocytes was significantly reduced (Figure 1C). A more detailed assessment of T cell subsets showed that the proportion of immature CD4^+^CD8^+^ thymocytes decreased markedly (Figure 1D). The percentage of myeloid cells in BM and erythroid cells in BM and spleen remained stable (Supplementary Figure 1G-H). Within the bone marrow, there was an immediate decrease in the percentages of B lymphoid cells, while all the other subsets remained unchanged (Supplementary Figure 1I-K). Further analysis indicated that the decrease in B lymphoid cells was most profound among pro-B/pre-B and immature B cells (Figure 1E).

**Figure 1.**
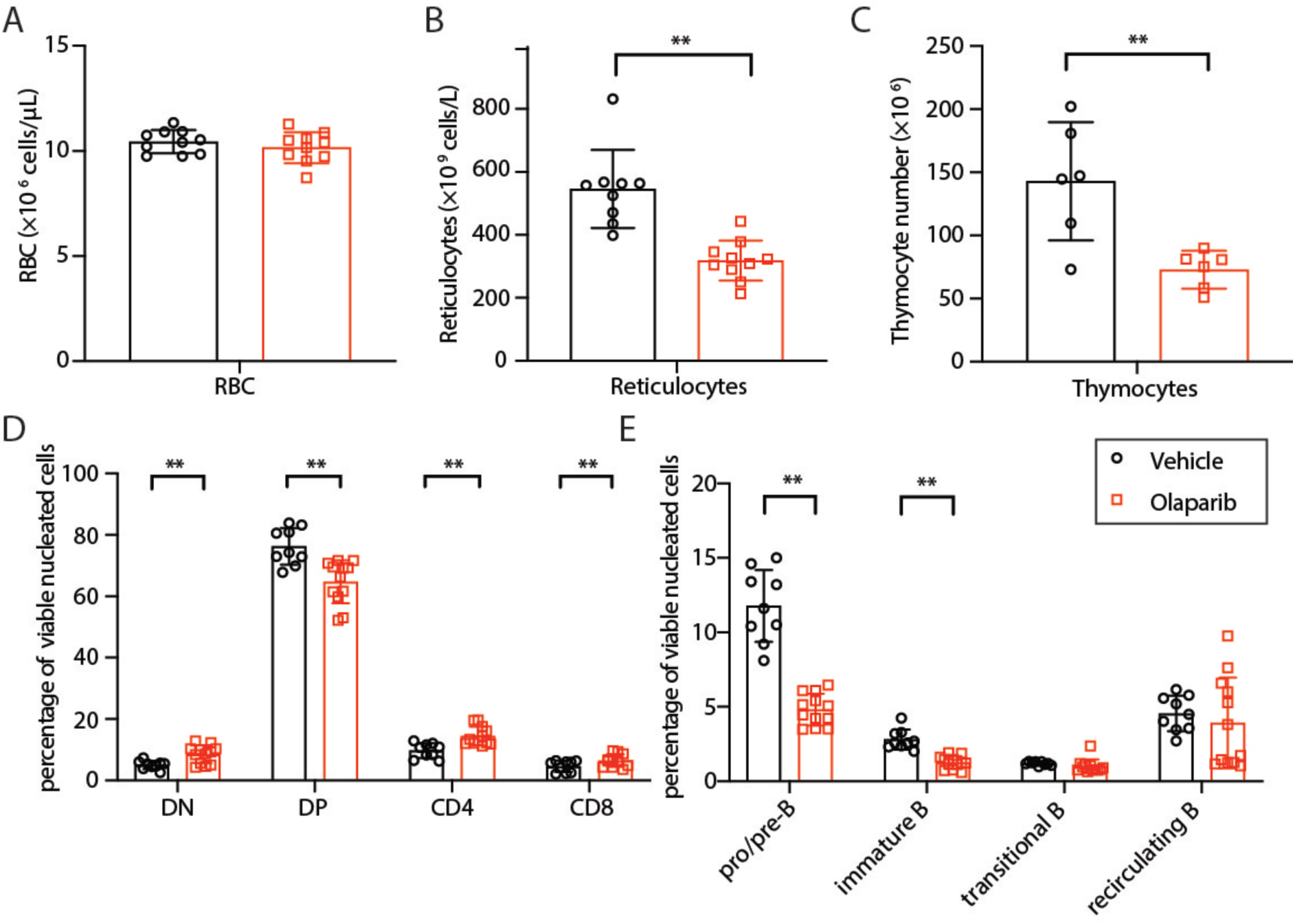
Acute impacts of olaparib treatment in C57BL/6 mice. Red blood cells (RBC) counts in peripheral blood (A), reticulocyte counts in peripheral blood (B), thymocyte counts (C), proportion of CD4 and CD8 positive cells in the thymus (D) and pro/pre-B and immature B percentages in BM (E) in C57BL/6 mice after 4 days of treatment with olaparib or vehicle. The data shown represent means ± 1 S.D. derived from vehicle-treated (n=9) or olaparib-treated (n=12) mice. P values were calculated using an unpaired two-tailed Student’s t test, * p < 0.05, ** p < 0.01.

To monitor the toxicity over a longer period, we extended olaparib treatment to 21 days. The changes observed were largely consistent with the 4 day treatment. The RBC count stayed the same, although the reticulocyte number was significantly lower (Supplementary Figure 2A-B). However, the drop was not as robust as that seen after 4 days of treatment (∼25% vs ∼42%). Similar to short term treatment, the counts of thymocytes, particularly those of the immature CD4^+^CD8^+^ thymocytes, and the counts of pro-B/pre-B and immature B cells in BM decreased substantially (Supplementary Figure 2C-E). Unlike short-term PARPi treatment, treatment for three weeks did induce a drop in myeloid cells, both in the marrow and in peripheral blood (Supplementary Figure 2F).

### A DNA repair focused CRISPR/Cas9 screen revealed loss of CHK2 blunts PARPi response in *Eµ-Myc* pre-B lymphoma cells

To determine whether we could recapitulate the sensitivity of immature B cells to PARPi *in vitro*, we tested the response of primary pro-B/pre-B lymphoid cells from C57BL/6 mice cultured on OP9 cells. We found that pro-B and pre-B subsets grown from the BM were highly sensitive to killing by olaparib (EC_50_= 97±42 nM) (Figure 2A, Supplementary Figure 3A). Cell lines established from *Eµ-Myc* pre-B lymphomas (31) were also readily killed by olaparib (EC_50_ = 227±12 nM) (Figure 2B, Supplementary Figure 3B), although they exhibited a higher IC_50_ than primary cells. The availability of blood cell lines sensitive to PARPi provided an opportunity to explore genetic factors that modulate drug response. Specifically, we sought to identify a strategy to prevent treatment related hematological toxicity.

**Figure 2.**
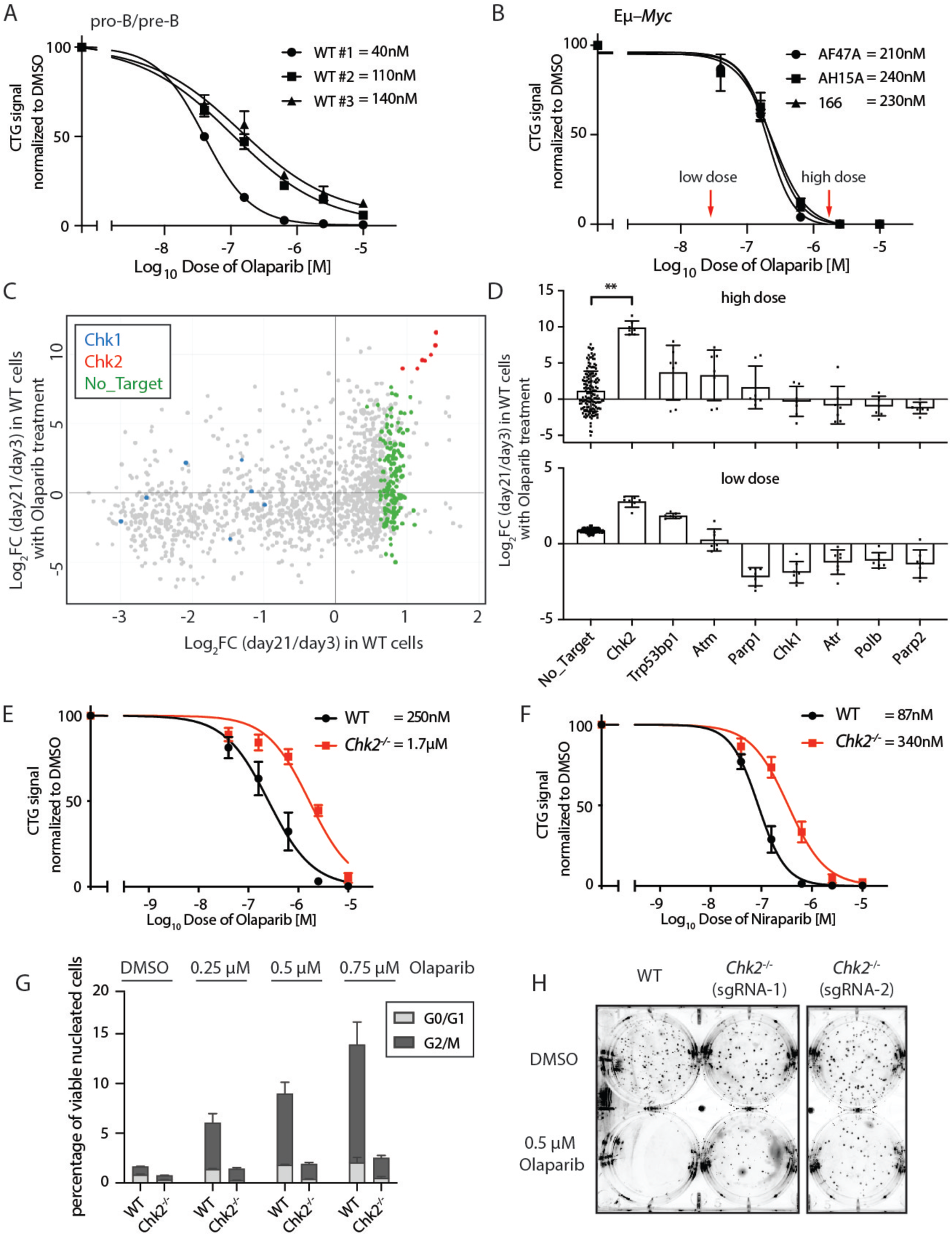
Loss of CHK2 alleviates PARPi induced cytotoxicity in *Eµ-Myc* lymphoma cells. Primary BM pro-B/pre-B cells (A) or Eµ-Myc lymphoma cells (B) were treated with olaparib (0-10µM) for 48 hours. Cell viability was measured using CellTiter-Glo and results expressed relative to untreated cells. EC_50_ values are reported. Red arrows indicate olaparib doses used for the CRISPR/Cas9 screens (40nM and 2µM). (C) CRISPR/Cas9 screening results. We plotted the log_2_ fold change of the relative abundance of each sgRNA between day 3 days and day 21, for untreated cells (x-axis) and olaparib treated cells (y-axis). Select sgRNA are highlighted: *Chk2* (in red), *Chk1* (in blue) or non-targeting (in green). (D) Log_2_ fold changes of the relative abundance of sgRNA against *Parp1, Parp2, Polb* and components of the p53 pathway at day 21 *vs* day 3 in high (upper) or low (lower) doses of olaparib. (E-F) Impact of loss of *Chk2* on the sensitivity of *Eµ-Myc* lymphoma cells to the PARP inhibitors olaparib (E) and niraparib (F). (G) Cell cycle analysis of WT *Eµ-Myc* lymphoma cells and isogenic *Chk2*^*-/-*^ derivatives treated with olaparib (0, 0.25, 0.5 or 0.75 µM). (H) Impact of loss of *Chk2* on the long-term clonogenic survival of *Eµ-Myc* lymphoma cells treated with DMSO or 0.5 µM olaparib. BM pro-B/pre-B cells were sorted from three WT mice (A), three *Eµ-Myc* lymphoma lines (AF47A, AH15A, 166) (B), three isogenic Chk2-/- knockout derivates were used in (E-F), that were made from two parental *Eµ-Myc* lymphoma lines (2 in AF47A, 1 in AH15A). Data shown in (A-B, E-F) are means ± 1 S.D. at 48 hours from three independent experiments performed in triplicate. Data shown in (G) are means ± 1 S.D. from three independent experiments. Image shown in (H) is a representative image from three replicates.

We focused our attention on the DNA damage response and generated a custom CRISPR/Cas9 library to target established DNA repair genes. Our CRISPR/Cas9 library contained 7 sgRNA per gene, targeting 174 DNA repair genes, 10 essential genes (e.g. *Rpa3, Rpl11, Psmb2*) (32) and *Hprt1*, together with 150 non-targeting control sgRNAs. Cas9 expressing *Eµ-Myc* lymphoma cells were transduced with the sgRNA pools before treatment with olaparib over 21 days. Cells were treated with low dose olaparib (IC_10_: 40 nM) to identify synthetic lethality interactions, or high doses (IC_99_: 2 µM) to identify sgRNAs that confer resistance. An untreated control was included to identify genes that impact on cell growth or survival. sgRNA against essential genes, such as *Rpa3, Rpl11, Psmb2, Plk1*, were depleted over the 3 weeks of culture (Supplementary Figure 3C). While many studies have defined synthetic lethal interactions between PARPi and BRCA1/BRCA2, this was not evident in *Eµ-Myc* lymphoma cells. Guides targeting *Brca1* and *Brca2* were depleted in both olaparib treated cells and the untreated controls. We did find robust negative selection of multiple guides targeting *Polb* and *Parp2* upon treatment with olaparib (Supplementary Figure 3D). POLB has previously been reported as a synthetic lethal interaction with PARPi in mouse embryonic fibroblasts (33) and our results suggests this is conserved across tissues. The depletion of Parp2 sgRNAs suggests that simultaneous inhibition of both PARP1 and PARP2 leads to greater cell killing. This is consistent with prior observations that olaparib preferentially targets PARP1 (34-36). These results confirmed that the DNA repair CRISPR/Cas9 library provided a way to identify genes that modify olaparib response.

We next looked for genes mediating response. Culturing cells in high doses of olaparib (IC_99_: 2 µM) resulted in strong selection of guides targeting the serine threonine kinase *Chk2*. This enrichment was observed with all 7 guides targeting *Chk2*, but the degree of enrichment varied between 490- to 2900-fold. When we looked at low dose olaparib treatment, it was obvious that *Chk2* guides were also enriched under these conditions. We did not see robust changes in sgRNAs targeting other p53 pathway components, such as *Chk1, Atm* or *Atr* (Figure 2D), but there was some enrichment for guides targeting *Tp53bp1*, which has been implicated in PARPi resistance (37-39). These results suggest some level of specificity for *Chk2* inhibition, within the broader p53 pathway, but they must be interpreted with caution, as some targets, like *Chk1*, were negatively selected in all conditions.

To verify our screen results, we generated isogenic *Chk2* knockout *Eµ-Myc* lymphoma derived cell lines using CRISPR/Cas9 and confirmed CHK2 loss attenuated olaparib-induced killing and cell cyle arrest (Figure 2E & 2G, Supplementary Figure 3E). Loss of CHK2 also conferred resistance to another PARPi, niraparib (Figure 2F), which is more efficient at trapping PARP on DNA (40). Increased tolerance to olaparib was also observed in long term clonogenic assays over 13 days (Figure 2H). Targeting CHK2 also impaired PARPi response in Hoxa9-Meis1 AML cell lines (Supplementary Figure 3F-G), indicating that the resistance phenotype is observed in myeloid cells. As part of our analysis we looked in greater detail at how blood cells respond to PARPi. We found that PARPi induces apoptosis in *Eµ-Myc* lymphoma cells, that can be blocked by combined loss of the effector proteins Bak and Bax (Supplementary 4A-B). Loss of CHK2 did not generally impair the apoptotic response, as these cells remained highly sensitive to the MCL-1 inhibitor (S63845) (41) (Supplementary Figure 3H). In keeping with prior work (30), we confirmed that CHK2 deficient cells were more resistant to DNA damaging agents, like etoposide, but remained sensitive to direct activation of p53 through addition of an MDM2 inhibitor (RG-7388) (Supplementary Figure 3I-J). These findings fit with the general view that CHK2 is an important amplifier of DNA damage signalling via activation of p53.

### Activation of p53 mediates PARPi toxicity in normal blood cells, both *in vitro* and *in vivo*

CHK2 plays a critical role in regulating the activity of p53 in response to DNA damage. This occurs through a broader signalling network, in which DNA damage activates key regulatory kinases, like ATM, that activate CHK2, which then directly phosphorylates p53 and promotes its stabilisation (21). Stabilisation of p53 results in activation of downstream target genes that trigger cell cycle arrest and apoptosis (Figure 3A). p53 protein levels increased in response to treatment with olaparib in *Eµ-Myc* lymphoma cells, and this relies in part on CHK2, because the response was markedly delayed in *Chk2*^-/-^ cells (Figure 3B). Disrupting p53 also conferred resistance to both olaparib and niraparib (Figure 3C-D), suggesting that the immediate toxicity of PARPi in blood cells is mediated through activation of the p53 pathway. As expected, loss of p53 also limited killing by etoposide and an MDM2 inhibitor, but did not prevent killing by an MCL-1 inhibitor (S63845) (Figure 3E-G).

**Figure 3.**
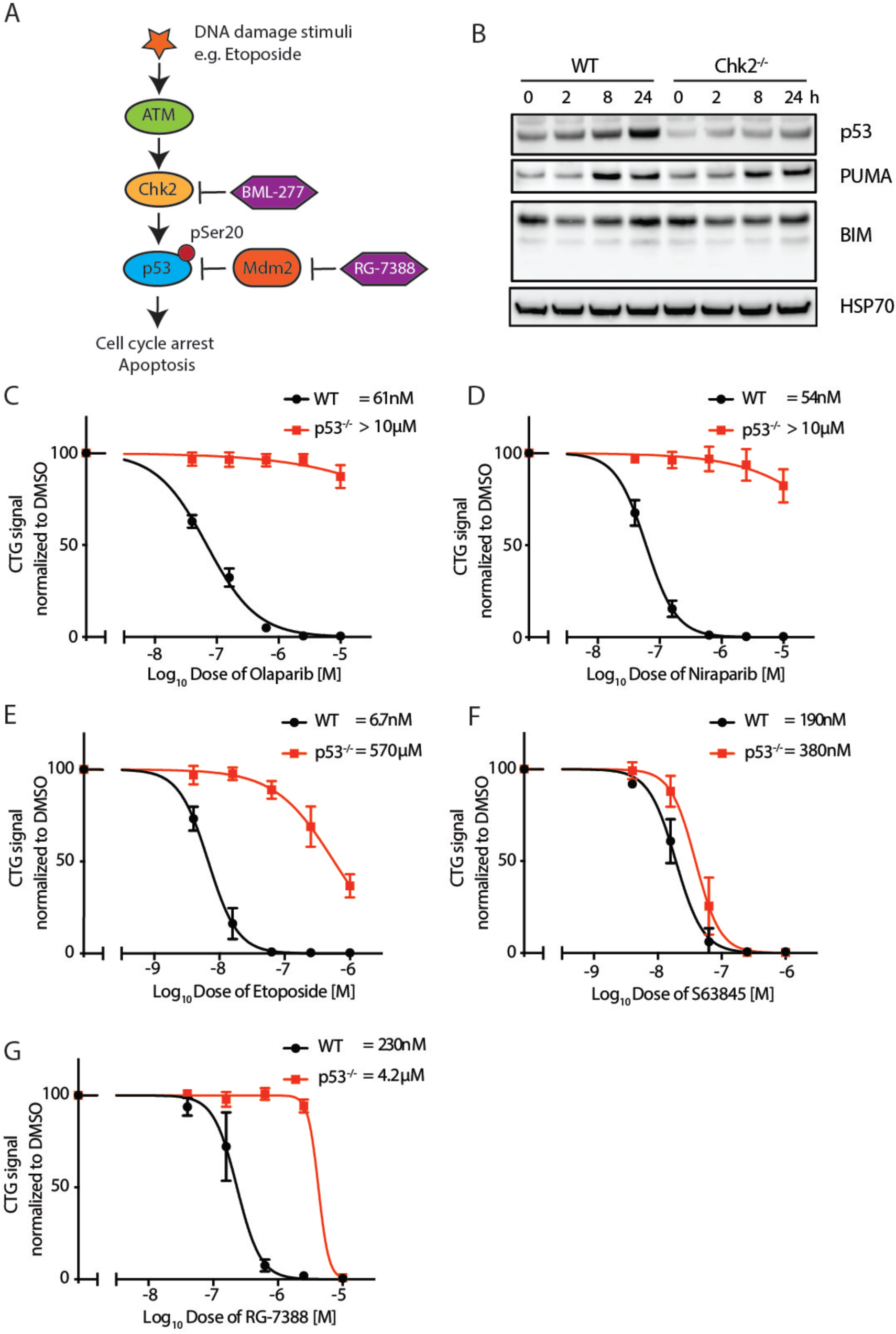
CHK2-dependent p53 activation is crucial for PARPi induced cytotoxicity. (A) Model of CHK2-dependent p53 activation. In response to DNA damage, the DNA damage sensor ATM is activated. ATM activates CHK2, and CHK2, in turn, phosphorylates p53 at Serine 20, resulting in its stabilisation and activation of target genes. Inhibitors used to manipulate the activity of the pathway in this study include the CHK2i, BML-277, and an MDM2 inhibitor, RG-7388. (B) Immunoblots for p53, BIM and PUMA in WT and *Chk2*^-/-^ *Eµ-Myc* lymphoma cells treated with 1 µM olaparib for 0-24 hours in the presence of 25 μM Q-VD-OPh (caspase inhibitor). HSP70 serves as a loading control. (C-G) Cell titre glow assays were used to assess the impact of loss of p53 on the sensitivity of *Eµ-Myc* lymphoma cells to olaparib (C), niraparib (D), etoposide (E), S63845 (F) and RG-7388 (G), following 48 hours of drug treatment. Data shown in (C-G) are the mean ± 1 S.D. from three independent experiments using two *Eµ-Myc* lymphoma lines, and their derivatives, performed in triplicate.

To evaluate whether p53 is crucial for olaparib induced toxicity on hematopoietic cells *in vivo*, we assessed olaparib response in *Trp53*^-/-^ mice (C57BL/6 background). The absence of p53 prevented the loss of reticulocytes and reduction in thymus cellularity typically seen with short term olaparib treatment (Figure 4A-B). Loss of p53 was also sufficient to correct the defects observed in the T and B cell lineages - specifically we did not see a change in the representation of CD4^+^CD8^+^ T cells in the thymus, or changes in B cell progenitors in the bone marrow (Figure 4C-D). These results confirmed that p53 plays a critical role in mediating the acute toxic effect of olaparib.

**Figure 4.**
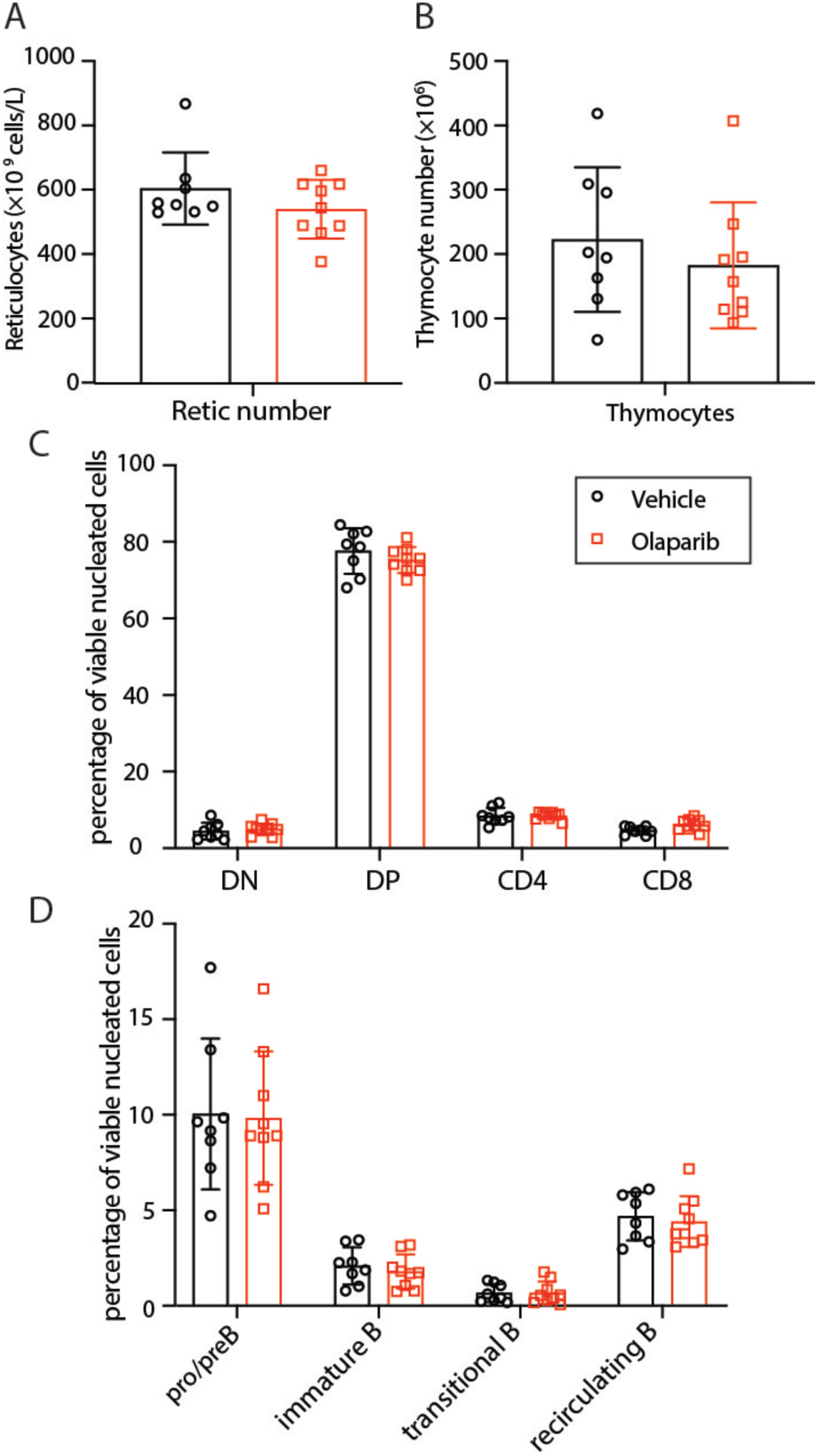
Olaparib treatment fails to reduce reticulocyte and BM immature B cell counts in *p53*^-/-^ mice. Reticulocyte counts in the peripheral blood (A), thymocyte counts (B), proportion of CD4 and CD8 positive cells in the thymus (C), and proportion of pro-B/pre-B and immature B percentages in BM (D) in *p53*^-/-^ mice after 4 days of treatment with olaparib or vehicle. The data shown in (A-D) represent means ± 1 S.D. derived from vehicle-treated (n=8) or olaparib-treated (n=9) *p53*^-/-^ mice. No significant differences were observed between the treatment arms.

### Ser23 phosphorylation on p53 is essential for PARPi induced cytotoxicity in *Eµ-Myc* lymphoma cells

Given that CHK2 regulates p53 activity via direct phosphorylation, we sought to determine whether blocking p53 phosphorylation would impair PARPi induced cell killing. To do this we used a CRISPR/Cas9 base editor to target a key site in p53 that is phosphorylated by CHK2, which is Ser23 in mice (thought to be equivalent to Ser20 in humans). The base editor was expressed in *Eµ-Myc* lymphoma cells and introduced a series of mutations in and around Ser23, which included synonymous, non-synonymous and nonsense mutations (Supplementary Figure 5A), which could be monitored by small amplicon sequencing. The editing was inefficient, with edits observed in ∼8% of cells in the pool, with a diverse set of mutant alleles. We treated the pool of modified cells with different drugs and monitored mutant allele frequencies.

We first looked at the behaviour of nonsense alleles in p53. We found that treatment with an MDM2 inhibitor or a PARP inhibitor resulted in enrichment of nonsense alleles in p53 (Figure 5 and Supplementary Figure 5B, light and dark blue bars). For example, in the presence of the MDM2 inhibitor RG-7388, we saw a 65-fold increase in the abundance of the c.54-55 CC>TT p.Q19* allele over three days, and a 108-fold enrichment over 7 days, with this allele ultimately contributing >25% of the total population. Nonsense alleles were not enriched upon treatment with the MCL-1 inhibitor, where p53 is not required for an effective response at high doses (Figure 3F). Synonymous mutations in p53 were found at consistent allele frequencies across all treatment conditions, with some mild depletion when conditions favoured outgrowth of p53-deficient cells (Figure 5, light and dark green bars).

**Figure 5.**
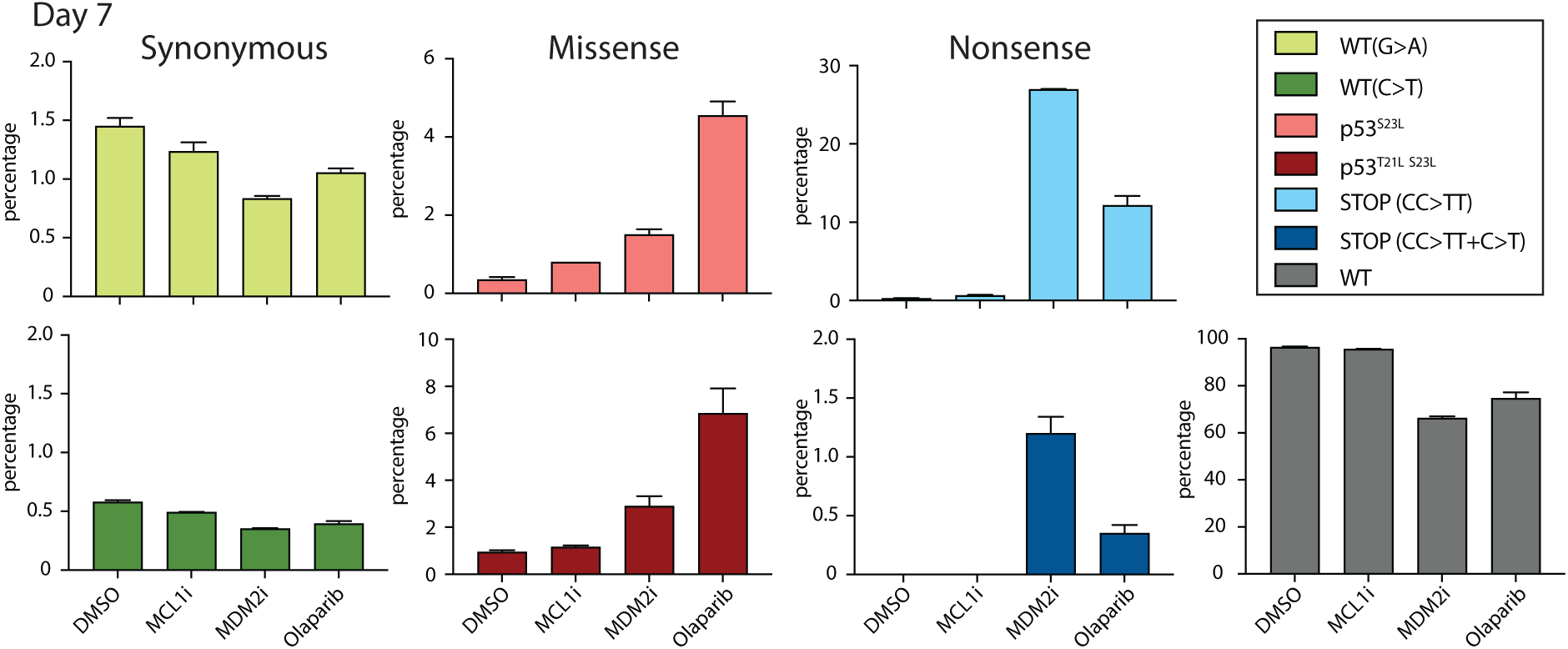
p53 Ser23 phosphorylation contributes to PARPi induced cytotoxicity. A base editor was used to introduce mutations in key phosphorylation sites for p53. The representation of each edited allele was tracked before and after treatment with an MCL1 inhibitor (S63845), MDM2 inhibitor (RG-7388), olaparib or a DMSO control, for 7 days. Results are shown for these alleles: WT (G>A) (light green), WT (C>T) (dark green), S23L (pink), T21I+S23L (red), STOP (CC>TT) (light blue), STOP (CC>TT+C>T) (dark blue) and WT (not edited) (grey). Some alleles were grouped, because they had the same impact on the protein (detailed in Supplementary Figure 5A). Data shown were means ± 1 S.D. from three independent cultures for each condition generated from the same editing event. Data obtained at three days are shown in Supplementary Figure 5B.

Our editing approach yielded four mutant alleles where p53 Ser23 had been mutated to leucine (p53^S23L^), including two with a concurrent change at Thr21 (p53^T21L S23L^). We found enrichment for mutant alleles encoding p53^S23L^ and p53^T21L S23L^ after treatment with either the MDM2 inhibitor or olaparib. In the presence of olaparib, mutating p53 Ser23 alone resulted in a 8.7-fold enrichment over three days (Supplementary Figure 5B), and a 13-fold enrichment after 7 days (Figure 5, pink bars) and consistent results were seen with p53^T21L S23L^ (Figure 5, red bars). Although the differences were modest, both p53 missense mutants showed stronger enrichment in olaparib than the MDM2 inhibitor, which was the opposite trend to that seen with the nonsense mutants. Failure to strongly enrich the p53 Ser23 missense mutants in the presence of the MDM2 inhibitor suggests they retain functional activity. These results support the view that phosphorylation of p53 at Ser23 is required for optimal activation of p53 in response to olaparib.

### CHK2i (BML-277) antagonized PARPi in p53 proficient pro-B/pre-B cells, but not in ovarian cancer cells

We next investigated whether pharmacological inhibition of CHK2 could also attenuate PARPi induced cytotoxicity. Here we used a CHK2 specific inhibitor BML-277 (42, 43), which can effectively block olaparib induced CHK2 phosphorylation (Supplementary Figure 6). Addition of BML-277 lessened PARPi induced cytotoxicity in primary pro-B/pre-B cells and in *Eµ-Myc* lymphoma cells (Figure 6A,C). The antagonist influence of BML-277 was not observed in *Chk2*^-/-^ or *p53*^-/-^ cells, although in both cases the killing by olaparib was far less effective than in wildtype cells (Figure 6B, D, E). Looking at intermediate doses, we observed that BML-277 at 0.64 and 2.5 µM can robustly inhibit cytotoxicity of olaparib in WT cells, but this was not seen with *Chk2*^-/-^ or *p53*^-/-^ cells (Figure 6F). We used a Bliss summary index to assess the interaction across all tested conditions. Olaparib and BML-277 showed an antagonist effect in p53 proficient pro-B/pre-B and *Eµ-Myc* lymphoma cells, but not in *p53*^*-/-*^ or *Chk2*^-/-^ cells, where there was some mild synergy (Figure 6H).

**Figure 6.**
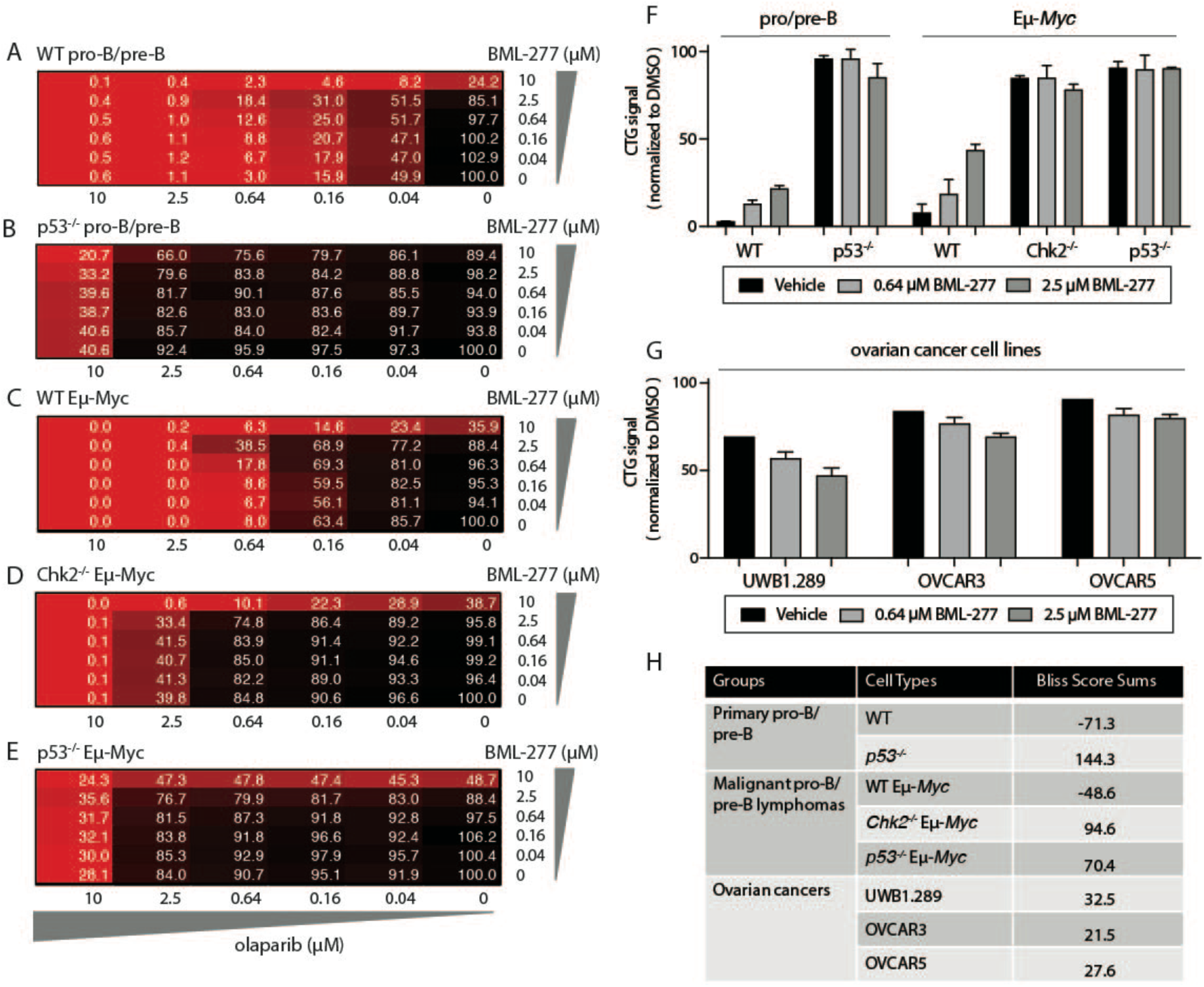
PARPi induced cytotoxicity is antagonised by a Chk2 inhibitor in p53 proficient pro-B/pre-B cells, but not in p53 deficient ovarian cancer cells. (A-E) Olaparib and BML-277 combination treatment in WT (A) and p53^-/-^(B) BM primary pro/pre-B cells, and in WT (C), *Chk2*^-/-^ (D) and *p53*^-/-^ (E) *Eµ-Myc* lymphoma cells. Cell titre glow assays were performed after 48 hours of treatment, with values expressed relative to the DMSO control. Drug concentrations are indicated on x and y axes of each matrix, and results from a representative experiment have been shown. Values from the triplicate experiments are shown for select doses in (F). (F-G) Cell viability in WT and *p53*^-/-^ BM pro/pre-B cells, and WT, *Chk2*^-/-^ and *p53*^-/-^ Eµ-*Myc* lymphoma cells (F), or in the ovarian cancer cell lines UWB1.289, OVCAR3 or OVCAR5 (G), treated with 0.64 µM olaparib together with DMSO or BML-277 (0.64 or 2.5 µM). Viability was assessed using CellTiter-Glo assay and results are normalised to the DMSO control. (H) A summary of Bliss scores obtained for the combination of olaparib and BML-277 in normal and malignant blood cells and ovarian cancer cells. All the primary data for pro-B/pre-B cells shown in (A-B, F, H) are derived from cells isolated from three independent WT or *p53*^-/-^ mice, each with three replicates; All the Eµ-*Myc* data shown in (C-E, F, H) are measured at 48 hours from two independent WT *Eµ-Myc* lymphoma lines and isogenic *p53* or *Chk2* knockout derivatives, each with three replicates; data from the ovarian cancer cell in (G-H) are measured at 96h from three independent experiments, each with two replicates.

Given that PARPi are typically applied in p53-deficient cancers, we did not think that concurrent CHK2 inhibition would negatively influence their anti-cancer activity; however, we decided to test this interaction using ovarian cancer cell lines. *TP53* is mutated in >96% of high grade serous ovarian cancers (24, 44). Here we tested the interaction of olaparib and BML-277 in three p53 deficient ovarian cancer cell lines: UWB1.289 (BRCA1 mutant), OVCAR3 (BRCA proficient) and OVCAR5 (BRCA proficient) (45, 46). Olaparib does not induce rapid killing in these cell lines, so we extended the length of treatment to 96 hours to assess its influence on cell proliferation. BML-277 did not block olaparib induced cytotoxicity in any of these cell lines (Figure 6G, H and Supplementary Figure 7A-C). This suggests that the use of BML-277 should not negatively affect olaparib efficacy.

## Discussion

PARPi are being pursued for the treatment of many different cancers. While it is clear that cells with defective HR are exquisitely sensitive to PARPi, these agents have a widespread impact on DNA repair, as well as other cellular processes. By assessing the impact of PARPi on the normal murine hematopoietic system, we found that reticulocytes, immature B cells and immature CD4^+^CD8^+^ thymocytes are acutely sensitive, but longer-term treatment also impacts myeloid cells (Figure 1 & Supplementary Figure 2). These results mirror findings from *Parp2* knockout models (19, 20, 47), indicating the toxic impact of these inhibitors are from on-target inhibition. They are also consistent with the observation that hematological toxicity is a common side effect of PARPi treatment, with anemia and neutropenia being common adverse events (17). We found that PARPi rapidly induced both cell death and cell cycle arrest in blood cells, and that these acute effects relied on activation of the p53 pathway. Here we outline a strategy to dampen the p53 pathway to reduce the toxic influence of PARPi on the blood.

We developed a DNA repair CRISPR/Cas9 library to investigate ways to modulate PARPi response in blood cells. As a model, we used *Eµ-Myc* lymphoma cells, the malignant counterpart of BM pro-B/pre-B cells, that are highly responsive to PARPi (Figure 2B, Supplementary Figure 3B). Upon treatment with PARPi we observed depletion of sgRNAs targeting genes that are established synthetic lethal partners, namely *Polb, Parp2* and Fanconi anemia genes (40, 48), but this did not include *Brca1* or *Brca2*, which were depleted in both treated and untreated cells. The depletion of *Brca1* and *Brca2* guides suggests a strong reliance on the HR pathway in these cells, which is not surprising given that they divide rapidly and retain a functional p53 response. Nonetheless, these results gave us confidence that we could reliably detect genetic determinants of PARPi response.

We focused on the genes that were required to potentiate the cytotoxic activity of PARPi. We found that guides targeting *Trp53bp1* and *Chk2* were enriched in *Eµ-Myc* lymphoma cells upon PARPi treatment (Figure 2D). Subsequent validation work with CHK2 confirmed this observation held true with other PARPi, like niraparib (Figure 2F), and was also seen in myeloid cells (Supplementary Figure 3F-G). CHK2 acts as a crucial bridge between the DNA damage sensing machinery and the p53 pathway. CHK2 directly phosphorylates p53, and we found that loss of p53 was sufficient to block cell killing by PARPi, both *in vitro* and *in vivo* (Figure 3-4). To more directly assess the importance of the phosphorylation of p53 by CHK2, we used a cytosine base-editor to mutate key phosphorylation sites, specifically Ser23, which is thought to be equivalent to human Ser20 (21). p53 phospho-site mutants carrying Ser23Leu, were enriched in the presence of PARPi, and this enrichment was less obvious when we used an MDM2 inhibitor to stabilise p53 protein. This implies a degree of specificity for the DNA damage response. *Chk2* knockout mice are viable and healthy (23), so we went on to investigate therapeutic targeting of CHK2 as a way to limit the toxic impact of PARPi on the blood.

The idea of targeting p53 or CHK2 for radio- or chemo-protection has been investigated for decades. More recent work has highlighted that this strategy may only have utility for select chemotherapeutic drugs. A screening approach found that loss of CHK2 protected blood cells from topoisomerase II inhibitors, like etoposide and doxorubicin, but not from topoisomerase I inhibitors, where, perhaps counter-intuitively, it seemed to sensitise (30). Our findings extend this concept to include protection from PARPi. One concern that we had in pursuing efforts to blunt PARPi activity was that we would compromise their anti-cancer activity. It is, however, important to consider that PARPi are almost always applied in p53-deficient cancers. For example, PARPi have been most successful in high-grade serous ovarian cancers, where p53 is mutated in nearly all cases (24, 44), offering a reminder that the efficacy of these agents does not require a functional p53 pathway. We demonstrated that CHK2 inhibitors blunt the response of hematopoietic cells to PARPi treatment, but did not influence growth inhibition of ovarian cancer cell lines (Figure 6). If these results are replicated *in vivo*, then co-treatment with a CHK2 inhibitor could offer a way to limit the toxic influence of PARPi on healthy cells.

Importantly, others have suggested that combined inhibition of CHK2 and PARP may be a way to potentiate the killing of cancer cells, presumably by influencing cell cycle progression via a p53-independent mechanism (26, 27). There are several factors that could account for these somewhat contradictory conclusions. First, we note that we saw PARPi antagonism at low concentrations of the CHK2 inhibitor BML-277, but saw enhanced killing at the highest concentration. By examining drug responses in different genetic backgrounds, including CHK2- and p53-deficient cells, we found that the antagonism at low doses of BML-277 relied on the presence of CHK2, but that synergy at high doses did not. Therefore, it is possible that the assumed synergy is a result of off-target activity. Another factor that could account for differences is the level of specificity of the CHK2 inhibitor, as our genetic studies suggest any inhibition of CHK1 could have a strong anti-proliferative effect. Indeed, work in another Myc-driven lymphoma model used a dual CHK1/2 inhibitor and found synergistic killing with PARPi (26). This observation requires further investigation, as we predict very different outcomes from treatments that target CHK1, which are being pursued as potential combination therapies for PARPi, along with other agents that target the DNA damage response and cell cycle checkpoints, such as ATR and WEE1 inhibitors (49). Lastly, it is possible that the functional interaction between CHK2 and PARPi may differ based on the cell type, or on the underlying tumor genotype. Our *in vivo* treatment model and work with primary B cell cultures supports the view that a selective CHK2 inhibitor would antagonise PARPi function.

Our results support the view that targeting CHK2 may offer some protection against hematopoietic toxicity due to PARPi treatment. There are risks associated with dampening the p53 pathway in normal tissues. Genetic association studies have linked germline loss-of-function alleles in *CHK2* to increased cancer risk (50); however, the increase in this risk appears modest, and may only apply in select cancer types (51). CHK2-deficient mice are viable and healthy (23), which suggests that transient inhibition of CHK2 should be tolerated, but it remains to be seen whether sustained inhibition of the pathway is a viable approach, particularly in the context of other anti-cancer treatments. More promising is recent work showing that that pharmacological targeting of CHK2, even transiently, can block the toxic influence of radiotherapy and chemotherapy on oocytes (28, 29), which is being pursued as a way to protect fertility during cancer treatment. Our results suggest that this protective influence extends to other tissues and treatments. The development and application of PARPi in HR deficient cancers represents an important conceptual step for cancer medicine, because it represents a shift towards treatments that target tumor-specific dependencies and vulnerabilities. Yet these drugs have a broad impact on multiple DNA repair pathways, as well as many other cellular processes (52). Dampening the p53 pathway, through inhibition of CHK2, offers a way to exacerbate the difference in response between HR deficient cancers and HR proficient normal tissues, which should broaden the therapeutic window for PARPi. This could have important implications for those patients that experience severe and persistent haematologic toxicity.

## Materials and methods

### Cell lines and mice

Hoxa9-Meis1 acute myeloid leukemia (53) and Eμ-*Myc* lymphoma (54) cell lines were generated and cultured as previously described. Human ovarian carcinoma cell lines UWB1.289, OVCAR3 and OVCAR5 were originally purchased from ATCC. UWB1.289 cells were cultured in 50% RPMI-1640 Medium (Gibco, Catalog No. 61870127) and 50% MEGM (Mammary Epithelial Growth Medium (Lonza MEGM Bullet Kit, CC-3150) supplemented with 3% Fetal Bovine Serum (FBS, Sigma, 10099-141), grown in 5% CO_2_ atmosphere at 37 °C. OVCAR3 and OVCAR5 were cultured in RPMI-1640 Medium with 10% FBS in 5% CO_2_ atmosphere at 37 °C.

### *In vivo* olaparib treatment

Mice were maintained on a C57BL/6 background. WT mice at 6-8 weeks were used for drug treatments. p53^-/-^ mice less than 10 weeks old were used for treatment and any with thymomas were excluded. For *in vivo* olaparib treatment, the vehicle was 10% DMSO/10% 2-hydroxypropyl-β-cyclodextrin (Sigma, H-107) in PBS. Olaparib (MedChemExpress, HY-10162) was dissolved in DMSO to 100 mg/ml and stored at - 20°C. One hour before use, an aliquot was thawed and diluted 1:10 with vehicle. Mice were treated with vehicle or olaparib (150 mg/kg) by intraperitoneal injection for four consecutive days, or for three weeks (5 days on, 2 days off). Experiments were conducted according to the guidelines of the Walter and Eliza Hall Institute Animal Ethics Committee (2016.023, 2018.046).

### Hematopoietic cell subset staining

Organs were harvested from mice 24 hours after the last injection. Cell suspensions were prepared and resuspended in 10mL Iscove’s Modified Dulbecco’s Medium (IMDM, Thermo Fisher, 12440053). 1×10^5^ APC labelled counting beads (BD, 340487) were added to 100 µL of cells to perform a cell count. BM cells were stained with B220-APC (clone RA36B2), IgD-PE (clone 11-26c.2a) and IgM-FITC (clone 5-1). BM, spleen and LN cells were stained with B220-FITC (clone RA36B2), CD4-PE (clone GK1.5), CD8-APC (clone 53-6.7), Mac1-PerCP.Cy5.5 (clone M1/70) and Gr1-PerCP.Cy7 (clone RB6-8C5). Thymocytes were stained with CD4-PE, CD8-APC. BM and spleen cells were stained with CD71-FITC (clone R17217.1.3) and TER119-PE (clone TER-119).

### CRISPR/Cas9 gene knockout

The lentiviral-based CRISPR/Cas9 editing system has been described (55). Briefly, two independent sgRNA against mouse *Chk2* (GTATACATAGAGGATCACAG and GCTGGAGACAGTGTCTACCC) or mouse *Trp53* (56) were cloned into pKLV-U6gRNA(BbsI)-PGKpuro2ABFP (Addgene, 50946). Lentiviral packaging plasmids (1µg pCMV-Rev, 1.5µg pMDLg/pRRE, 1µg pCMV-VSV-G) were co-transfected with a Cas9 lentiviral vector (56), or sgRNA plasmids (1.5µg), into HEK293T cells in a 10cm dish using 24µL FuGENE HD Transfection Reagent (Promega, E2311). Lentiviral supernatants were harvested 48h after transfection and filtered through a 0.22µM filter (Millipore, SLGV033RS). To generate Cas9 expressing cells, 1mL of Cas9 viral supernatant was added to 1×10^6^ cells growing in a 6-well plate plate in 3mL of complete media with polybrene (4mg/mL) (Sigma, H9268). The cell-virus mixture was centrifuged at 3000rpm at 25°C for 1h followed by incubation at 37°C overnight. The next day cells were washed and re-suspended in fresh media without polybrene. After a minimum of three days in culture, viable mCherry positive cells were sorted on AriaIII cell sorter (BD) the expanded for one week. Cells were then transduced with sgRNA virus following the same protocol and single mCherry/BFP double positive cells were sorted into 96-well plates. Sequencing of genomic DNA from clonal cell lines using the Illumina MiSeq sequencing platform (San Diego, CA, USA) has been described (56).

### CRISPR/Cas9 base editing

To generate a base editor plasmid with fluorescence selectable marker, hA3A-BE3 element were amplified from pCMV-hA3A-BE3 plasmid (Addgene, 113410) with the primers sets (BE3_PCR_F and BE3-PCR_R) using Pfu DNA polymerase (Promega, M7741). Amplified fragments were digested with XhoI and NotI and cloned into pCAGIG plasmid (Addgene, 11159) which was linearized using the same restriction enzymes to generate pCAGIG-hA3A-BE3.

BE3_PCR_F gaattcctcgagagagccgccaccatggaagccagcc

BE3_PCR_R acgtgcggccgcttagactttcctcttcttcttggg

For the mouse p53 Ser23 edit, *Eµ-Myc* cells were transduced with sgRNA lentivirus targeting an adjacent region (AGGAGACATTTTCAGGCTTA). After a minimum of three days in culture, viable BFP positive cells were sorted on Aria-III cell sorter (BD) and expanded for one week. pCAGIG-hA3A-BE3 plasmid was delivered into cells using SF Cell Line 4D-Nucleofector X Kit (Lonza, #V4XC-2032) using programme CA-137. Viable GFP/BFP double positive cells were sorted 36h after nucleofection.

Cells were then treated with DMSO, 500µM S63845 (Active Biochem, A-6044), 1µM RG-7388 (CHEMGOOD, C-1287) or 2µM olaparib (Selleckchem, S1060) for up to 7 days. Cells were taken at day 3 and day 7 for DNA extraction, followed by sequencing of the p53 Ser23 adjacent regions.

### CRISPR/Cas9 screens

A custom pooled DNA repair lentiviral sgRNA library in the backbone pRSG16-U6-sg-HTS6C -UbiC-TagRFP-2A-Puro was synthesised by Cellecta Inc. To prepare the library lentivirus, 1.5µg of the library plasmid was used for co-transfection for each 10cm dish of HEK293T using the same method outlined above. A total of ten dishes of HEK293T cells were used and the supernatant was combined, filtered through a 0.22 µM filter and aliquoted.

Two independent Cas9-GFP expressing Eµ-*Myc* cell lines were used for sgRNA library lentiviral transduction. For each line, 1×10^7^ cells were transduced and a transduction efficiency of ∼20% was achieved (fold representation for each sgRNA >1,200). Puromycin (1 mg/ml) was added to the cells 48h post transduction and maintained for 3 days to select for sgRNA expressing cells. 5×10^6^ cells (fold representation for each sgRNA >3,000) cells were collected and frozen down for a baseline control (day 3). The remaining cells were split between three conditions; they were treated with 40nM or 2µM olaparib, or left untreated. 5×10^6^ cells were collected every 3 days starting from day 12 to 21.

Genomic DNA was extracted from these cell pellets using DNeasy Blood & Tissue Kits (Qiagen, 69506). Libraries were prepared using the NGS Prep Kit for sgRNA Libraries in pRSG16/17 (KOHGW, CELLECTA, LNGS-120) and Supplementary Primer Set for LNGS-120 (CELLECTA, LNGS-120-SP) following the user manual and sequenced on Illumina NextSeq platform. Data was analysed using Model-based Analysis of Genome-wide CRISPR-Cas9 Knockout (MAGeCK) package (57).

### Immunoblotting and antibodies

Immunoblotting was performed as previously described (58). Transferred membranes were probed with mouse anti-HSP70 (clone N6, a gift from Dr Robin Anderson, Peter MacCallum Cancer Centre, Melbourne), mouse anti-p53 (clone PAb 122, ThermoFisher, MA5-12453), mouse anti-CHK2 (clone 1C12, Cell Signaling, 3440), a polyclonal rabbit anti-BIM (Stressgen, AAP-330-E), rabbit anti-PUMA (ProSci, 3043), or rabbit anti-phospho-CHK2 (Ser516) (Cell Signaling, 2669).

### BM pro-B/pre-B cell staining, sorting and culture

OP9 stromal cells were cultured in alpha-MEM medium containing 10% FBS, 1 mM L-glutamine, 1mM HEPES, 1mM Sodium Pyruvate, 50 μM β-mercaptoethanol, 100U Penicillin and 100µg/ml Streptomycin. Bone marrow from WT or *p53*^-/-^ mice was stained with B220-APC (clone RA36B2), IgD-PE (clone 11-26c.2a) and IgM-FITC (clone 5-1). B220^+^IgD^-^IgM^-^ cells were sorted on an Aria-III cell sorter (BD), the cells were resuspended in IMDM medium with 10% FBS and 10ng/mL murine IL-7, and cultured on OP9 stromal cells for use in cell viability assays.

### Cell viability assays

For drug sensitivity assays, WT *Eµ-Myc* lymphoma cells or Hoxa9-Meis1 AML cells, or their derivatives, were seeded into 96-well plates at a density of 1×10^5^ cells/well and treated with serial diluted drugs (5-point 1:4 dilution), including olaparib, niraparib (Selleckchem, S2741), S63845, etoposide (Selleckchem, S1225), RG-7388. Cell death was assessed at 48 hours using propidium iodide (PI) (Sigma, P4864) uptake, measured on an LSRIIW flow cytometer (BD). For CellTiter-Glo assays, 1×10^4^ cells were seeded into white 96-well plate (Greiner, 655083) and cell viability was assessed using the CellTiter-Glo Luminescent Assay (Promega, G9241).

To test the interaction between olaparib and BML-277 (Selleckchem, S8632), primary pro-B/pre-B cells (1×10^4^), Eµ-*Myc* lymphoma cells (1×10^4^), or ovarian cancer cells (2×10^3^) were seeded into 96-well white plates and treated with both drugs, either individually, or in a combination matrix (each drug 5-point 1:4 dilution, olaparib horizontal dilution, BML-277 vertical dilution). Cell viability was determined at 48 hours (pro-B/pre-B or Eµ-*Myc*) or at 96 hours (ovarian cancer cell lines) using Cell Titer Glo Luminescent Assay. Bliss scores were calculated as previously described (59).

### Clonogenic assays

500 WT Eµ-*Myc* lymphoma cells, or their isogenic *Chk2*^-/-^ derivatives, were seeded in IMDM medium supplemented with 10% fetal bovine serum and 0.3% agarose in a 6-well plate. Cells were cultured at 37°C for 13 days in the presence of 0.5 µM olaparib or a DMSO control, then imaged on a ChemiDoc Touch Imaging System (Bio-Rad, 1708370).

### Cell cycle analysis

2×10^6^ WT Eµ-*Myc* lymphoma cells, or their isogenic *Chk2*^-/-^ derivatives, were treated with DMSO, or olaparib (0.25, 0.5 or 0.75 µM) for 24 hours, followed by incubation with 3 µg/mL BrdU (Sigma, 19-160) for 2.5 hours. Cells were collected and stained with BrdU-APC and 7-AAD using BD Pharmingen BrdU Flow Kit (BD, 552598) following manual instructions, followed by analysis using an LSR IIW (BD).

## Supporting information

Supplementary Figures 1-7 and Supplementary Table descriptions

Supplementary Table 1 - CRISPR Targets

Supplementary Table 2 - CRISPR Guide Counts

## Acknowledgments

The authors wish to acknowledge Andreas Strasser and Raelene Endersby for thoughtful feedback on the manuscript. We wish to thank Rachel Hancock, Stephanie Bound and Laura Dunleavy for assistance with animal husbandry and drug treatment experiments, Chris Riffkin and Ksenija Nesic provided guidance with ovarian cell lines, and Martin Pal and Marco Herold for advice with CRISPR base editing. This research has been supported by project funding from the National Health and Medical Research Council of Australia (Project Grant to IJM 1145912), the Cancer Council of Victoria (IJM) and the Stafford Fox Medical Research Foundation (CLS and CV). Fellowship support was provided from the Felton Bequest (IJM) and the Victorian Cancer Agency (Clinical Fellowship to CLS CRF16014, MCRF15018 to IJM), and scholarship support from the Leukaemia Foundation and the Haematology Society of Australia and New Zealand (EL). Research was also made possible through the Australian Cancer Research Foundation, Victorian State Government Operational Infrastructure Support and Australian Government NHMRC IRIISS. The funders had no influence over the final content of the manuscript.

