## Supplementary Figures 1-7 and Supplementary Table descriptions for "CHK2 inhibition provides a strategy to suppress hematological toxicity from PARP inhibitors"

Ian J. Majewski

The Walter and Eliza Hall Institute of Medical Research


##### Supplementary Figures and Legends (7)

S Figure 1. Hematological profiling following short term olaparib treatment in C57/BL6 mice.
S Figure 2. Hematological profiling following three weeks of olaparib treatment in C57/BL6 mice.
S Figure 3. Chk2 loss alleviates PARPi induced cytotoxicity in leukemia cell lines.
S Figure 4. Olaparib triggers Bax and Bak mediated apoptotic cell death in Eμ-*Myc* lymphoma cells.
S Figure 5. Details of the base substitutions induced in *Trp53* by the hA3A-BE3 base editor.
S Figure 6. CHK2 inhibition by BML-277 can prevent phosphorylation of p53 in response to olaparib.
S Figure 7. No antagonist effect between olaparib and BML-277 in *p53* deficient ovarian cancer cell lines.

##### Supplementary Table Descriptions (2)

S Table 1. Gene list for CRISPR/Cas9 DNA repair gene library.
S Table 2. Count table and fold enrichment for screens performed with the CRISPR/Cas9 DNA repair gene library.

### Supplementary Figures and Legends

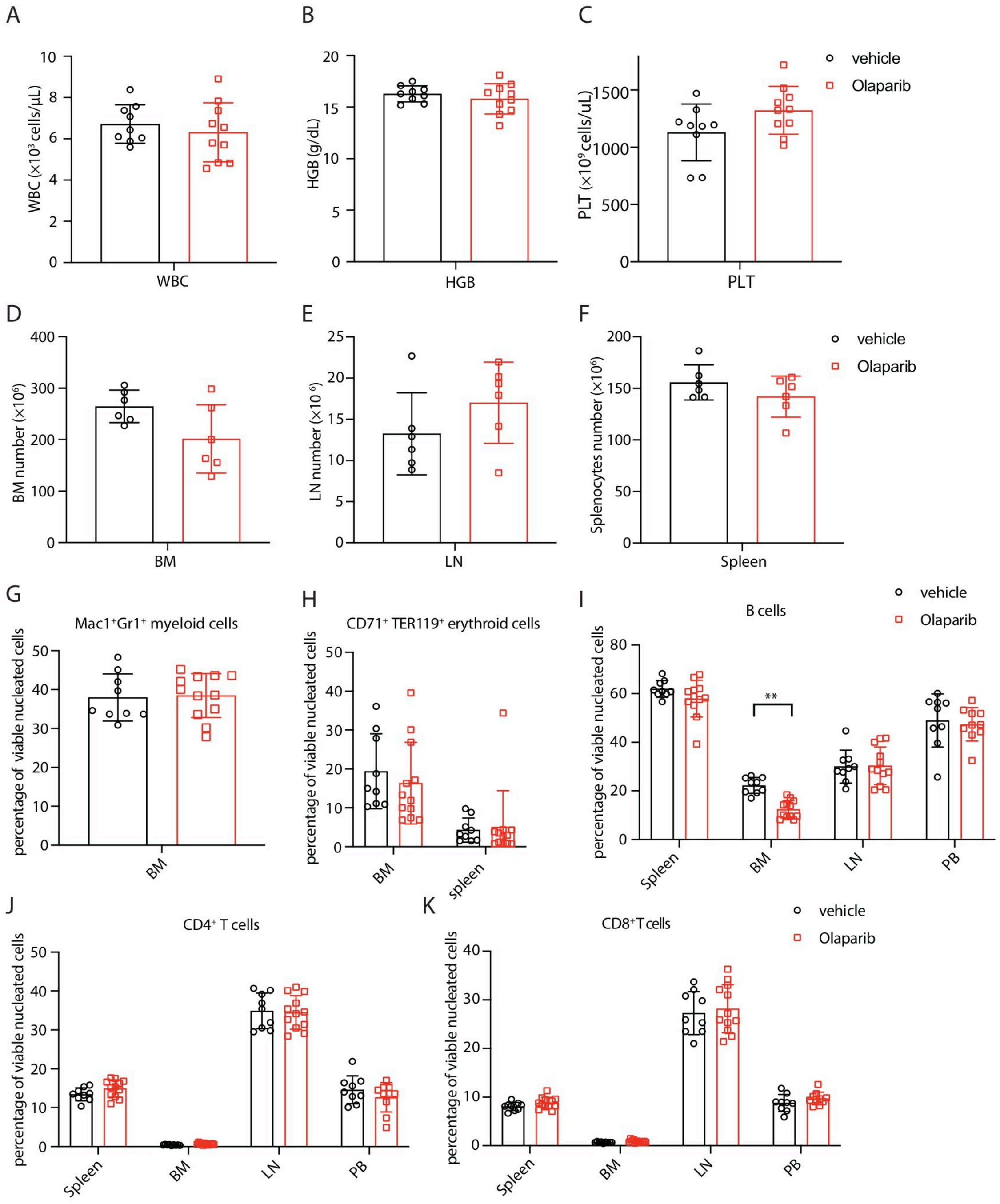


**Supplementary Figure 1. Hematological profiling following short term olaparib treatment in C57/BL6 mice.** (A-C) White blood cell counts (WBC) (A), hemoglobin (HGB) (B) and platelet counts (PLT) (C) in C57BL/6 mice treated with olaparib or a vehicle control for 4 days. BM (D), LN (E) and spleen (F) cellularity following olaparib treatment for 4 days, and flow cytometric assessment of myeloid cells in BM (G), erythroid cells in spleen or BM (H), and B cells (I), $CD4^+$ T cells (J) or $CD8^+$ T cells (K) in spleen, BM, LN or PB. Individual values are plotted from vehicle-treated (n=9) or olaparib-treated (n=12) mice gathered over 3 independent experiments, together with the mean ± 1 S.D.

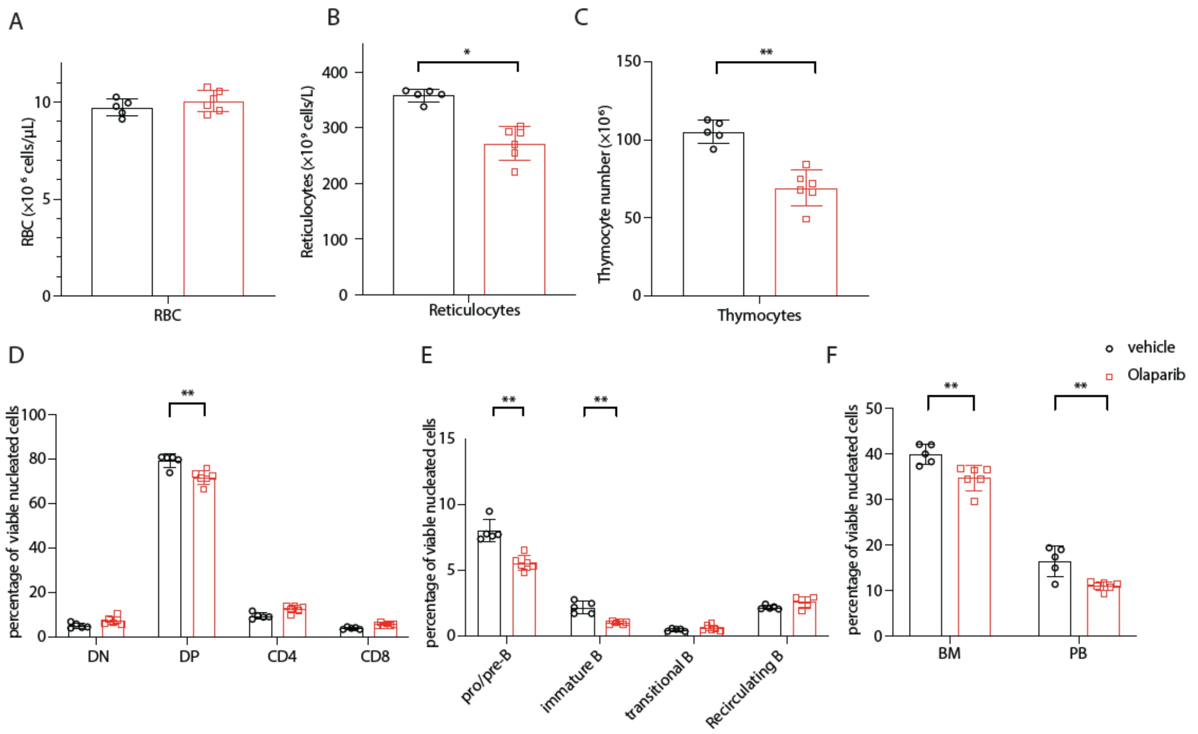


**Supplementary Figure 2. Hematological profiling following three weeks of olaparib treatment in C57/BL6 mice.** (A-F) RBC (A), reticulocyte counts (B), thymocyte counts (C), percentages of CD4 and CD8 positive cells in the thymus (D), percentages of BM pro-B/pre-B and immature B cells (E), percentages of BM and PB $Mac1^{+}Gr1^{+}$ myeloid cells (F) in C57BL/6 mice after 3 weeks of treatment with olaparib or vehicle. The data shown represent means ± 1 S.D. derived from vehicle-treated (n=5) or olaparib-treated (n=6) mice. P values were calculated using an unpaired two-tailed Student's t test, * $p < 0.05$, ** $p < 0.01$.

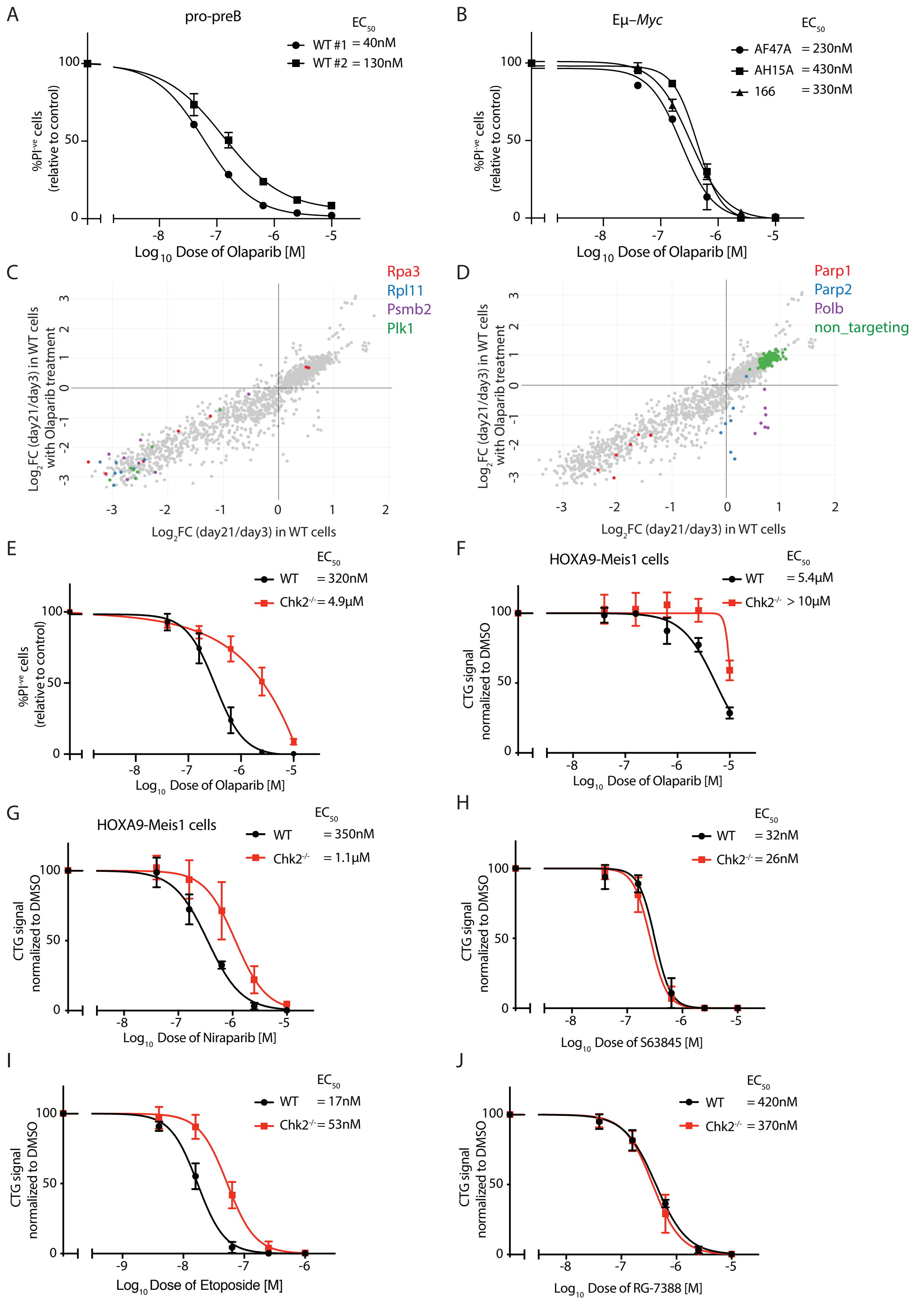


**Supplementary Figure 3. Chk2 loss alleviates PARPi induced cytotoxicity in leukemia cell lines.** Olaparib induced cell killing was measured by PI uptake in pro-B/pre-B cells (A, n=2) and in *Eμ-Myc* lymphoma cells (B, n=3), each with three replicates. (C-D) sgRNA abundance in *Eμ-Myc* lymphoma cells with or without olaparib (100 nM), showing the Log2 fold change in abundance between day 3 and day 21 of culture. (C) sgRNA targeting a select set of essential genes are highlighted (*Rpa3*, *Rpl11*, *Psmb2* or *Plk1*). (D) sgRNA targeting

*Parp1*, *Parp2* and *Polb* are highlighted, together with a set of non-targeting controls. (E) Cell viability was assessed with PI staining in wildtype and *Chk2*$^{-/-}$ *Eμ-Myc* lymphoma cells treated with olaparib. (F-G) CellTiter-Glo assays were used to assess the impact of loss of *Chk2* on drug sensitivities in HoxA9-Meis1 cells treated with olaparib (F) and niraparib (G). (H-J) CellTiter-Glo assays were performed with wildtype and *Chk2*$^{-/-}$ *Eμ-Myc* lymphoma cells treated with the MCL-1 inhibitor S63845 (H), etoposide (I) and RG-7388 (J).Data shown in (E-J) are means ± 1 S.D. at 48 hours from three independent experiments using two independent HoxA9-Meis1 or Eμ-*Myc* lines and isogenic *Chk2*$^{-/-}$ derivatives, each performed in triplicate.

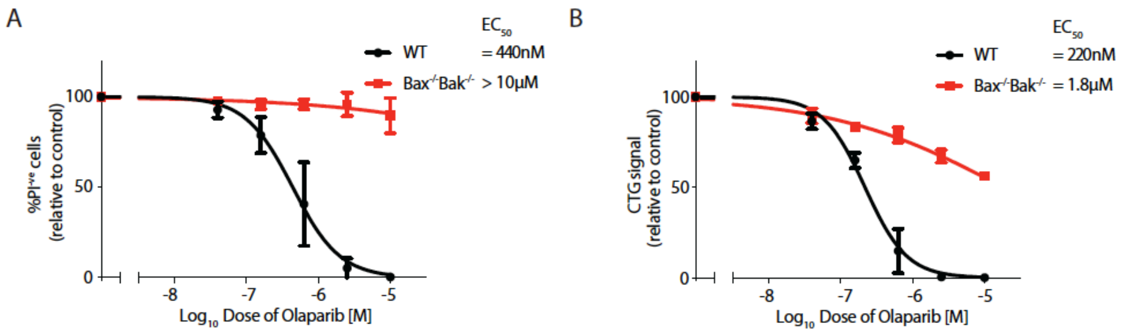


**Supplementary Figure 4. Olaparib triggers Bax and Bak mediated apoptotic cell death in Eμ-*Myc* lymphoma cells.** Olaparib response was assessed in WT and *Bax*$^{-/-}$*Bak*$^{-/-}$ *Eμ-Myc* lymphoma cells over 48 h. Cell viability was measured using PI staining (A), or with the CellTiter-Glo Assay (B). For both assays, values are shown relative to untreated controls. Data shown in (A-B) are means ± 1 S.D. from three independent experiments using two Eμ-*Myc* lines and isogenic *Bax*$^{-/-}$*Bak*$^{-/-}$ derivatives performed in triplicate.

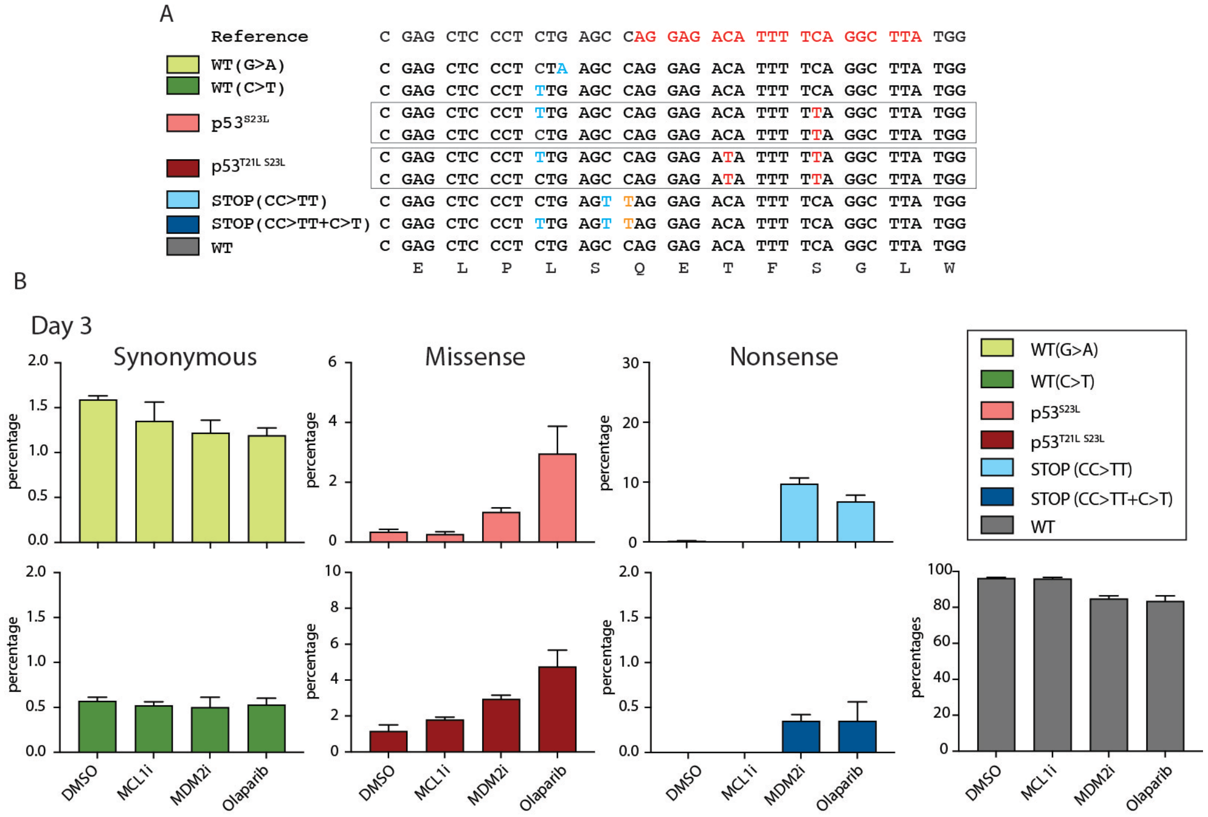


**Supplementary Figure 5. Details of the base substitutions induced in *Trp53* by the hA3A-BE3 base editor.** (A) The reference sequence for *Trp53* is shown at top, with the sgRNA sequence highlighted in red. Mutant alleles tracked in our study are shown. Base edits highlighted in blue indicate synonymous mutations, nucleotides in red indicate non-synonymous mutations, nucleotides in orange indicate nonsense mutations. Mutant alleles that are boxed together were grouped for quantification in the drug treatment experiments shown in Figure 5. (B) Quantification of edited alleles after 3 days of treatment with an MCL1 inhibitor (S63845), MDM2 inhibitor (RG-7388), olaparib or a DMSO control. Data shown were means ± 1 S.D. from three independent cultures for each condition generated from the same editing event. These cultures were continued for 7 days, with data presented in Figure 5.

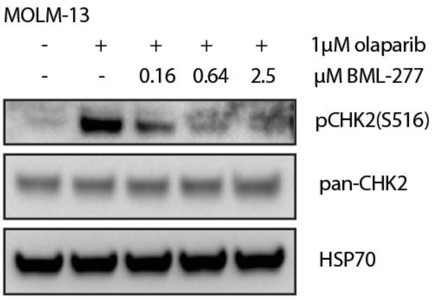


**Supplementary Figure 6. CHK2 inhibition by BML-277 can prevent phosphorylation of p53 in response to olaparib.** Immunoblots for phospho-CHK2 (Ser516) and pan-CHK2 in MOLM-13 leukemia cells treated with DMSO, 1μM olaparib or combined treatment with 1μM olaparib and BML-277 (0.16, 0.64 or 2.5 μM) for 4h. HSP70 serves as a loading control.

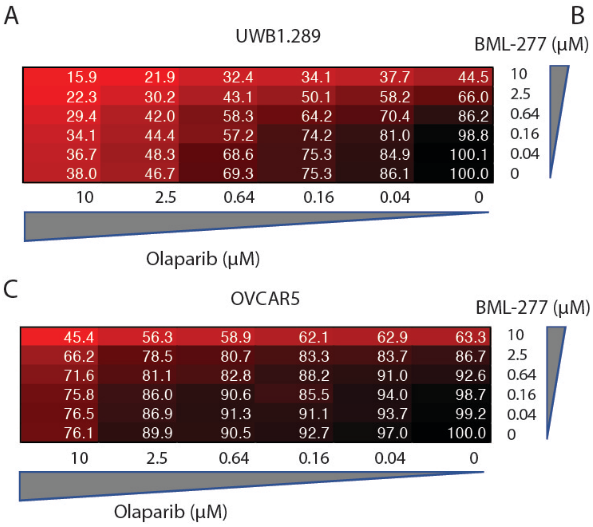


**Supplementary Figure 7. No antagonist effect between olaparib and BML-277 in *p53* deficient ovarian cancer cell lines.** Ovarian cancer cell lines UWB1.289 (A), OVCAR3 (B) or OVCAR5 (C) were treated with olaparib and BML-277 in combination for 96 h. Cell viability was measured using the Cell Titer-Glo assay and normalised to the control with DMSO treatment only. Data shown in (A-C) are representative of three independent experiments, each with two replicates. Results from the three independent experiments are shown for select doses in Figure 6H.

### Supplementary Tables

**Supplementary Table 1. Gene list for CRISPR/Cas9 DNA repair gene library.** We provide a list of the genes targeted in our CRISPR/Cas9 library design. The design includes 174 DNA repair genes, 10 essential genes, Hprt1 and non-targeting controls.

**Supplementary Table 2. Count table and fold enrichment for screens performed with the CRISPR/Cas9 DNA repair gene library.** Read counts and analysis of guide enrichment or depletion in cells treated with olaparib (40 nM or 2 μM) or an untreated control. We present guide counts in each condition, normalised counts and log fold changes.
